# Information processing by endoplasmic reticulum stress sensors

**DOI:** 10.1101/617217

**Authors:** Wylie Stroberg, Justin Eilertsen, Santiago Schnell

## Abstract

The unfolded protein response (UPR) is a collection of cellular feedback mechanisms that seek to maintain protein folding homeostasis in the endoplasmic reticulum (ER). When the ER is “stressed”, either through high protein folding demand or undersupply of chaperones and foldases, stress sensing proteins in the ER membrane initiate the UPR. Recently, experiments have indicated that these signaling molecules detect stress by being both sequestered by free chaperones and activated by free unfolded proteins. However, it remains unclear what advantage this bidirectional sensor control offers stressed cells. Here, we show that combining positive regulation of sensor activity by unfolded proteins with negative regulation by chaperones allows the sensor to make a more informative measurement of ER stress. The increase in the information capacity of the combined sensing mechanism stems from stretching of the active range of the sensor, at the cost of increased uncertainty due to the integration of multiple signals. These results provide a possible rationale for the evolution of the observed stress sensing mechanism.

## Background

The unfolded protein response (UPR) is a cellular stress response resulting from excessive accumulation of unfolded and misfolded protein in the endoplasmic reticulum (ER). Detection of heightened protein concentration within the ER lumen triggers accelerated protein folding and degradation within the ER along with decreased protein synthesis. If efforts to restore protein homeostasis are unsuccessful, the cell begins the process of apoptosis. Malfunction of the UPR has been implicated in numerous protein misfolding diseases (1), including type II diabetes mellitus (2; 3; 4) and neurodegenerative diseases (5). In yeast, the UPR is activated through a single pathway that depends on the transmembrane protein inositol-requiring enzyme 1 (Ire1) to transmit information about the activity of unfolded proteins within the ER lumen across the ER membrane. Oligomerization of Ire1 molecules activates its RNase domain, leading to the non-conventional spicing of HAC1 or (XBP1 in metazoan cells) mRNA (6). Spliced HAC1/XBP1 is translated to produce a bZIP transcription factor, which upregulates many genes related to protein homeostasis (7; 8). In higher eukaryotes, two additional additional branches of the UPR are regulated by the ER-membrane proteins protein-kinase-RNA-like endoplasmic reticulum kinase (Perk) and Activating transcription factor 6 (Atf6). While Atf6 signaling involves transport of the Atf6 out of the ER, Perk stress sensing functions analogously to Ire1 signaling, with the lumenal domain of Perk closely resembling that of Ire1 (9; 10; 11; 12). In this work, we focus on the stress sensing mechanisms of Ire1 and Perk.

Although significant progress has been made toward understanding the downstream cascade regulating chaperone production and ER-associated degradation (ERAD), the actual mechanism through which protein concentration in the ER is detected by the Ire1 and Perk luminal domains has remained controversial. Observations that the overexpression of the ER chaperone BiP reduced induction of the UPR lead many to believe that BiP sequestration by unfolded proteins triggered the stress response (13; 14). This idea was further bolstered by observations that a mutant partially-folded glycoprotein that could not bind BiP was also incapable of inducing the UPR (15), and that BiP constitutively binds Ire1, but shows a marked decrease in binding upon UPR activation (9; 16).

In contrast, there is growing evidence to support the notion of unfolded proteins acting as ligands through direct binding with Ire1 and Perk. For one, an Ire1 mutant lacking subregion V, which is thought to contain the BiP-binding domain, did not bind BiP, but had negligible effect on UPR activation compared to the wild type (17). Furthermore, a “core” mutant of Ire1, consisting of subregions II-IV, oligomerizes under physiological conditions, but does not activate the UPR unless ER stress is present (18). Ire1 dimers also display a shared groove similar to the peptide binding domain of major histocompatibility complexes (11), and mutations at the floor of this domain prevent binding of the model UPR-inducing misfolded protein CPY* in yeast (19; 20) and human cells (21). Similarly, the Perk lumenal domain selectively binds misfolded proteins through an analogous groove, which may then induce conformational changes that promotes oligomerization and kinase activity (22).

To integrate these observations, a hybrid model was proposed (23; 24), in which direct binding of unfolded proteins is necessary for full activation of the UPR, perhaps through the stabilization of Ire1 (and Perk) dimers, but the activation is buffered by competitive BiP binding to Ire1 and client unfolded proteins. Recent mathematical modeling of the UPR (25) has shown that controlling the activity of the response through the BiP-titration mechanism allows for a more efficient use of chaperone in mitigating unfolded protein stress in the ER, providing a rational for the evolution of BiP-modulated stress sensors. However, the same analysis demonstrated that including direct interactions between the sensors and unfolded proteins yielded no added benefit with regard to chaperone frugality. Hence, it remains unclear why the hybrid sensing mechanism that integrates signals of both chaperone and unfolded protein copy numbers has evolved.

In this work we hypothesize that combining both signals provides more information about the state of stress in the ER. To test this hypothesis, we construct a minimal mathematical model of the hybrid signaling network that contains the BiP-titration and direct-unfolded-protein signaling mechanisms as special cases. Using numerical and analytical techniques, we show that a BiP-mediated sensor can make a more informative measurement of available chaperone in the ER lumen by incorporating a measurement of the unfolded protein concentration directly. This advantage comes from extending the active range of the sensor response, at the cost of greater uncertainty due to integration of the unfolded protein-sensor interaction with the chaperone-sensor interaction.

## Model Description

To develop a model of stress signaling in the ER, we employ a modular approach and take advantage of the separation in timescales between the protein-protein interactions within the ER that determine the sensor activity, and the response time of the UPR. A schematic of the model is shown in Fig. 1 shows a schematic of the modular model, which consists of model for chaperone-assisted folding in the ER lumen (Folding module), and a stochastic model of the stress sensing proteins (Stochastic sensor module).

### Folding module

The first module describes the folding of unfolded proteins with the assistance of chaperones within the ER lumen. The chaperone-assisted folding is modeled as an enzymatic reaction

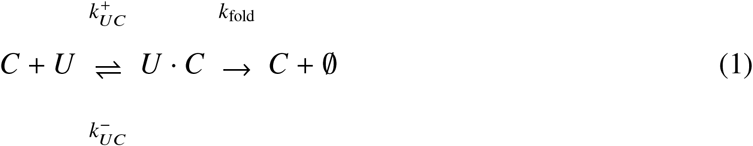

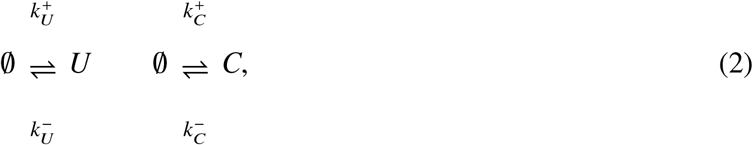

where *U* is an unfolded protein, *C* is a chaperone (e.g. BiP in metazoa), and *U* · *C* is a complex of chaperone and unfolded protein undergoing folding. Reaction (1) captures the binding of unfolded proteins by chaperones, and the subsequent folding and export of the folded protein from the ER. Reactions (2) represent influx of nascent proteins and chaperones into the ER, and their subsequent degradation or dilution. In general, binding of nascent proteins by chaperones occurs rapidly upon translocation into the ER (26), whereas folding of a nascent protein requires substantially more time (≳ 15 min) (27). The separation of these timescales allows for the approximation that the *U* and *C* are in a quasi-steady state with the folding complex *U* · *C* on timescales shorter than the folding time. Furthermore, we assume that significant changes in influx rate occur on a longer timescale than the *U* · *C* complex formation time so that, on the timescales of interest, the total copy number of unfolded protein in the ER, *U*_0_, and chaperone in the ER, *C*_0_, are conserved quantities. This assumption can be justified by the fact that the important timescale for signaling results from protein-protein interactions and is on the order of seconds, while significant changes in the influx rate require changes in gene transcription and protein translation, both of which vary more slowly. Hence, on the timescale of protein-protein interactions within the ER, the folding module reduces to a simple bimolecular reaction mechanism with conserved protein and chaperone copy numbers. This leads to (quasi) steady-state populations of unfolded proteins, chaperones, and folding complexes given by

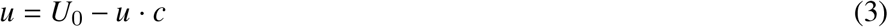

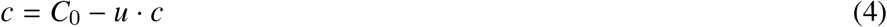

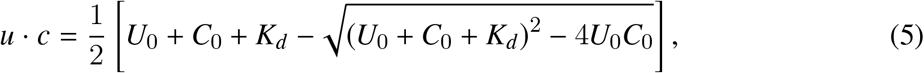

where *K*_*d*_ is the dissociation constant between unfolded proteins and chaperones, and *u*, *c*, and *u* · *c* are the steady-state copy numbers of unfolded proteins, unbound chaperons, and unfolded-protein-chaperone complexes in the ER, respectively. Without loss of generality, we set *K*_*d*_ = 1 and let all concentrations be measured relative to *K*_*d*_. Throughout our analysis, we will treat the populations of *u*, *c*, and *u* · *c* as deterministic, and focus on the fluctuations of sensor activity.

**Figure 1:**
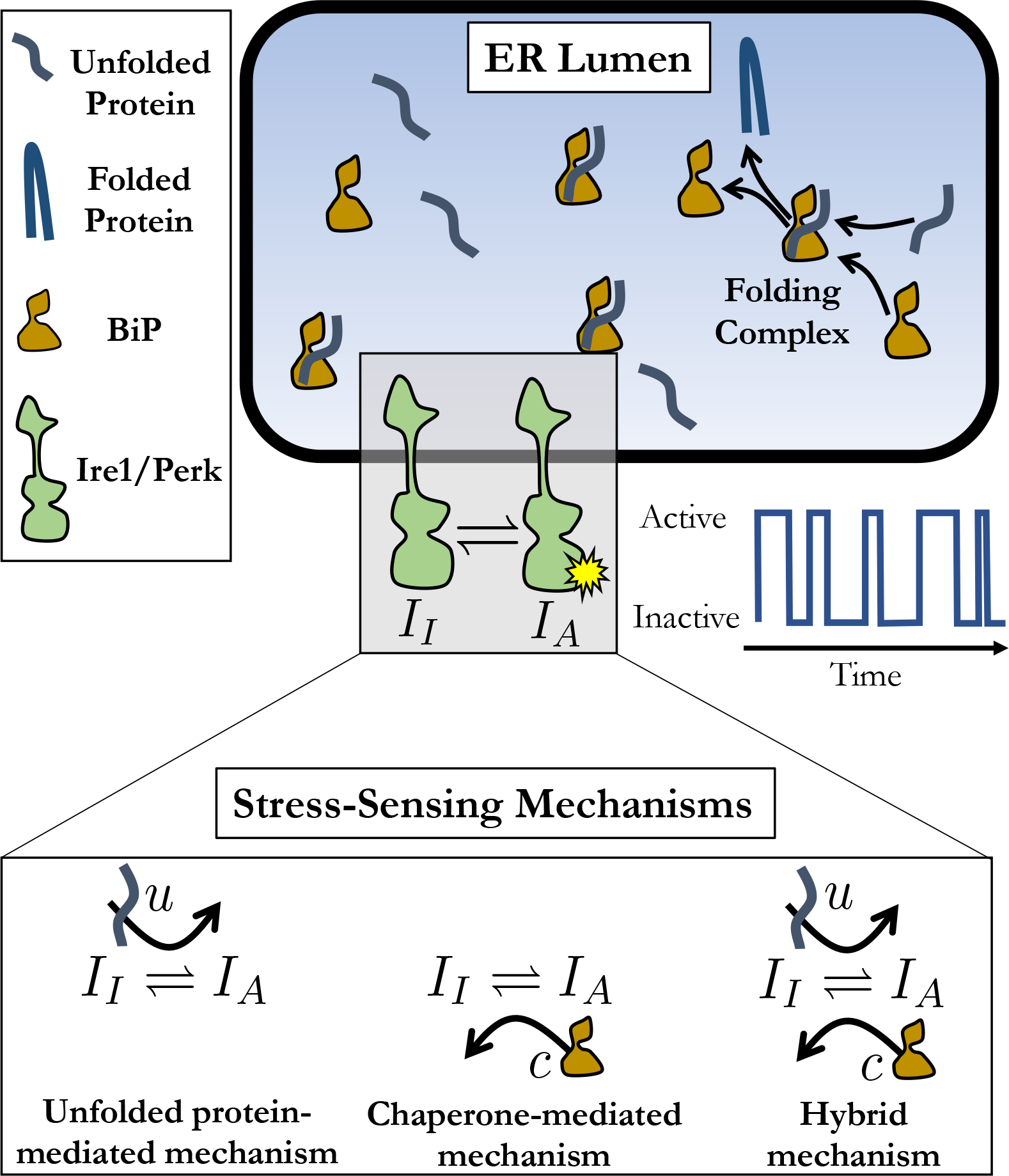
ER stress sensing model schematic. In the lumen of the ER, unfolded proteins are bound by the chaperone BiP, which aids folding. Eventually, the stable folded proteins are released from the folding complex, and exported from the ER. The folding process is monitored by the transmembrane stress-sensing proteins Ire1 and Perk. In our model, ER stress is measured through one of the three mechanisms depicted in the bottom panel, and transmitted as stochastic time series of the transmembrane sensor activation.

### Stochastic sensor module

The second module in our description of the ER stress sensing network is the transmembrane sensor. While metazoa have three distinct signaling pathways, we will focus here on the activation mechanisms of Ire1 and Perk, as they are thought to detect stress through the same mechanism and Ire1 is the most conserved of the three pathways. Our minimal model for the hybrid sensor mechanism consists of a two-state sensor with an activation rate that depends on the copy number of unfolded proteins and a deactivation rate that depends on the chaperone copy number:

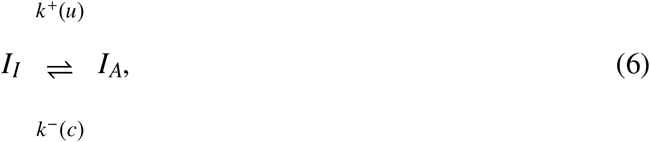

where *I*_*I*_ is the inactive state of the sensor, and *I*_*A*_ is the active state. Since the inputs *u* and *c* are assumed to change much more slowly than the time required for the sensors to probe the state of the ER, we are interested in the equilibrium fluctuations of the sensors. To this end, the equilibrium activation constant for the two state sensor is defined to be

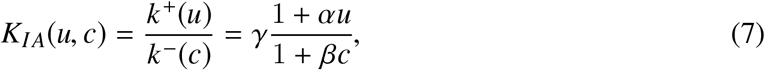

where *γ* controls the baseline scale of sensor activity, *α* dictates the sensitivity of the response to changes in unfolded protein copy number, and *β* sets the sensitivity of the response to changes in unbound chaperone copy number. In the limit as *α* → 0, *u* no longer directly influences the activity of the sensor, reducing to the mechanism in which sensor activity is regulated through chaperone titration only. On the other hand, in the limit as *β* → 0, the sensor activity is regulated only by direct interactions with unfolded proteins, with the chaperone providing no additional regulation of the sensor. Hence, this simple push-pull model for sensor activity encompasses the core regulatory mechanisms involved in Ire1 and Perk activity.

Taking *n*_*A*_ to be a random variable that represents the number of active sensor molecules (i.e., *I*_*A*_), and the total number of sensors in the system to be *N*_*I*_, the corresponding chemical master equations for reaction (6) is

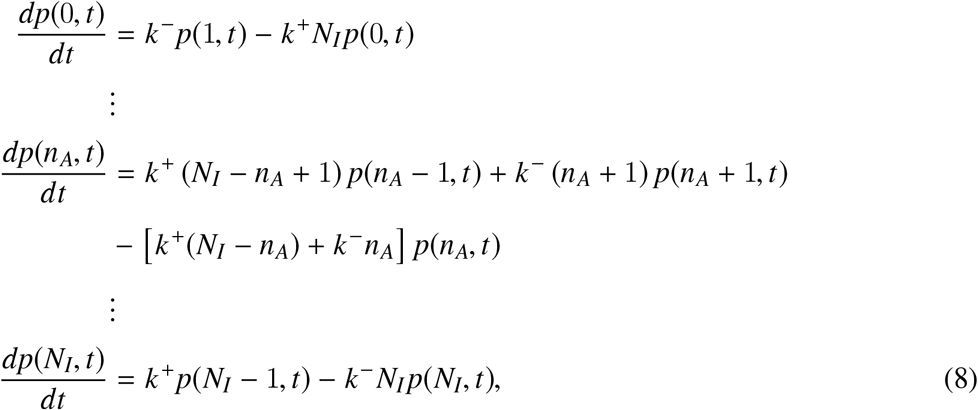

where *n*_*A*_ runs from 1 to *N*_*I*_ − 1. At steady state, equation (8) can be solved exactly (28), giving the probability of active sensors conditioned on the inputs (*u*, *c*):

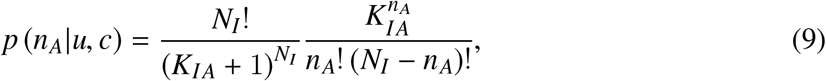

where the first term is a normalization constant that does not depend on *n*_*A*_, but does depend on *u* and *c* through *K*_*IA*_.

### Quantifying information capacity of ER stress sensors

In our model of ER stress signaling, there are two inputs to the system: (*U*_0_, *C*_0_). We refer to this set of variables as the “state” of the ER. Hence, the input (or prior) to our model is a joint distribution defining the state of the ER, *q*(*U*_0_, *C*_0_). However, the stress-sensing network is not necessarily seeking to measure either *U*_0_ or *C*_0_, but instead to measure the *stress*. While somewhat nebulously defined in the literature, ER stress should be a function of the state of the ER and quantify the potential for protein misfolding and aggregation. The simplest choice for a quantitative definition of ER stress would then be the number of unbound unfolded proteins, *u*(*U*_0_, *C*_0_), which we use here as our stress measure. Later, we will extend our analysis to consider the case in which the concentration of free chaperone, *c*(*U*_0_, *C*_0_) serves as the measure of ER stress.

We are interested in quantifying how well the sensor output *n*_*A*_ characterizes our ER stress measure. A common metric for quantifying the signal transduction quality in a sensory network is the mutual information between the input stimulus and the sensory response (29; 30; 31). For two distributions *X* and *Y* (e.g., input and output distributions), the mutual information between *X* and *Y* is given by

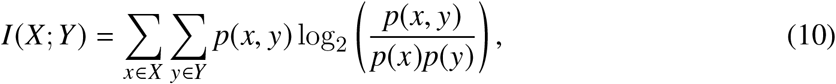

where *p*(*x*, *y*) is the joint probability distribution of *x* and *y* and *p*(*x*) and *p*(*y*) are the respective marginal probability distributions. Hence, our metric for sensor mechanism quality is *I*(*u*; *n*_*A*_), the mutual information between ER stress (which we initially take to be free unfolded protein concentration) and the output of the stress sensor.

To calculate the mutual information, it is necessary to assume a prior distribution for the input *q*(*U*_0_, *C*_0_). A common choice is to assume a uniform distribution over the input variables. However, in the case of ER stress, this leads to an unrealistically large range of possible values for free unfolded protein. Were the prior to be a uniform distribution in *U*_0_ and *C*_0_, the range of *u* would be from 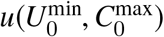 to 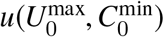. Yet, ER stress sensors are sensitive to departures of the system from homeostasis and will not necessarily need to measure the level of stress when the chaperone content is maximal and total protein load is minimal as this state is both unlikely to occur, and clearly not a state that requires a stress responses. Similarly, should the protein client load be exceptionally high and the chaperone copy number be at its baseline expression level, a response should have already been initiated. Hence, we must construct a more informed prior to draw more definitive conclusions about the quality of different stress sensing mechanisms. In particular, for the stress sensing mechanism to be responsive to stress at different processing capacities, it should retain a sensitivity to a given range of unfolded protein copy number as the copy number of chaperone changes. This ensures the homeostatic control mechanism is effective as the protein production capacity of the ER changes. The simplest prior distribution with this property is a uniform distribution in the copy number of unbound unfolded proteins, *u*, and in total chaperone copy number, *C*_0_,

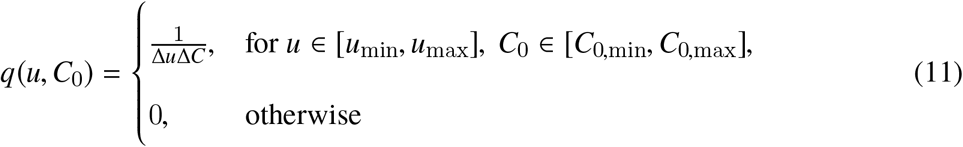

where Δ*u* = *u*_max_ − *u*_min_ and Δ*C* = *C*_0,max_−*C*_0,min_. Note that since *u* and *C*_0_ are typically quite large, we treat them as continuous random variables and *q*(*u*, *C*_0_) is a continuous distribution. *q*(*u*, *C*_0_) can readily be transformed into a distribution in terms of *U*_0_ and *C*_0_ by inverting equation (3), or into a distribution in term of unbound unfolded proteins and chaperons, *u* and *c*, using equation (4). With the prior distribution in this form, *q*(*u*, *c*), the probability distributions needed to calculate the mutual information in equation (10) can be readily calculated with the aid of the transfer function (9). Specifically, the marginal distribution of the input, *p*(*u*), is given by

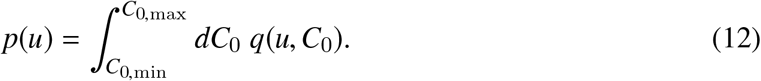

The joint distribution between unfolded protein and active sensors is given by

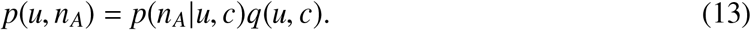

Lastly, the marginal distribution of the output is

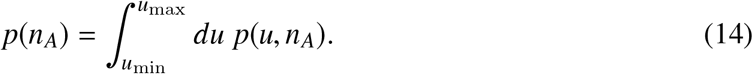

### Numerical calculation and optimization of mutual information

An exact, closed-form expression for the mutual information is challenging to obtain due to the difficulty in marginalizing the joint distribution of *u* and *n*_*A*_. Hence, we calculate the mutual information numerically. While the number of active sensors is already a discrete quantity, it is necessary to discretize the prior distribution to determine a numerical value for the mutual information. This is done by dividing the range of inputs into discrete bins such that there are *N*_*u*_ equally spaced bins of *u* values between *u*_min_ and *u*_max_, and *N*_*C*_ values of *C*_0_ between *C*_0,min_ and *C*_0,max_. The initial distribution is then transformed into a discrete distribution in the variables *u* and *c* with *N*_*c*_ bins between *c*_min_ and *c*_max_. The mutual information can then be computed by numerically calculating the necessary (discrete) probability distributions *p*(*u*), *p*(*n*_*A*_) and *p*(*u*, *n*_*A*_), and applying equation (10) directly.

In order the compare the information for different sensor mechanisms, it is pertinent to compare the maximal amounts of information each mechanism can transmit, i.e., the channel capacity, for across all sets of kinetic parameters. To calculate the channel capacity of each mechanism, the mutual information is numerically maximized over the kinetic parameters in each model: {*α*, *γ*} in the unfolded-protein mediated mechanism, {*β*, *γ*} in the chaperone-mediated mechanism, and {*α*, *β*, *γ*} in the hybrid mechanism. Details of the maximization procedure are provided in the Supplementary Material.

## Results

Using mutual information as a metric, we seek to quantify the effectiveness of different sensing mechanisms at monitoring protein homeostasis in the ER. In particular, we initially assume that the quantity of interest of the UPR regulatory network is the concentration of unfolded protein in the ER lumen and ask which sensing mechanism provides the most information about this quantity. Further, once we have determined which sensing mechanism is most informative, we would like to then understand how this mechanism is able to better measure the level of stress in the ER. We do this with a combination of computational and asymptotic techniques to discern how integrating the signals measuring unfolded protein and free chaperone can be most effectively achieved. Lastly, we ask how the optimal sensing mechanism changes if the quantity of interest (i.e. the measure of ER stress) is the concentration of free chaperone, as opposed to unfolded proteins.

### Direct sensing of unfolded proteins is more informative than chaperone-mediated sensing

Following optimization of the mechanism-specific rate parameters, the only unconstrained parameters in our model are the ranges of the input distributions, *C*_0 min_, *C*_0,max_, *u*_min_, *u*_max_, and the total number of sensors, *N*_*I*_. To show that the channel capacity of the direct sensing mechanism exceeds that of the indirect mechanism, we perform numerical calculations of the maximal mutual information for each mechanism across a broad range of prior distributions and for several values of *N*_*I*_ (see Supplementary Material). In Fig. 2, the maximal values of mutual information of each mechanism are displayed as heat maps for a ranges of prior distributions. In each case, *C*_0 min_ and *u*_min_ are fixed, and the upper bounds, *C*_0 max_ and *u*_max_ are varied over several orders of magnitude. Comparing the mutual information of the unfolded protein-mediated and chaperone-mediated mechanisms for any particular set of input parameters shows that the unfolded protein-mediated mechanism will always provide more information about the unfolded protein concentration in the ER than the chaperone-mediated mechanism. Quantitatively, the difference between the information between the unfolded protein- and chaperone-mediated mechanisms, Δ*I*_*u−c*_, ranges between 0.19-1.08 bits for the chosen set of prior distributions. Additionally, Fig. 2c shows the channel capacity for the hybrid mechanism. Since this mechanism has an additional free parameter compared to the unfolded protein- and chaperone-mediated mechanisms, it clearly can provide at least as much information as the better of the two special cases. Fig. 2c shows that, in general, the hybrid mechanism provides more information than either of the the other two mechanisms (Δ*I*_*h−u*_ ranges from 0.03-0.37 bits for chosen priors), and that this difference in greatest when the range of *u* is large and the range of *C*_0_ is small. Observing Fig. 2a shows the channel capacity for the direct mechanism is independent of the range of total chaperone in the input distribution. Intuitively, this makes sense as the direct mechanism only measures free unfolded protein concentration, and hence is decoupled from the chaperone concentration. This decoupling depends on our choice of input distributions: the probability distribution of free unfolded protein concentration between the chosen bounds are independent of chaperone concentration. This would not necessarily be the case for another choice of priors – for example, a uniform distribution of total unfolded protein (chaperone-bound and unbound) – in which case the free unfolded protein distribution would be coupled to the concentration of chaperone.

**Figure 2:**
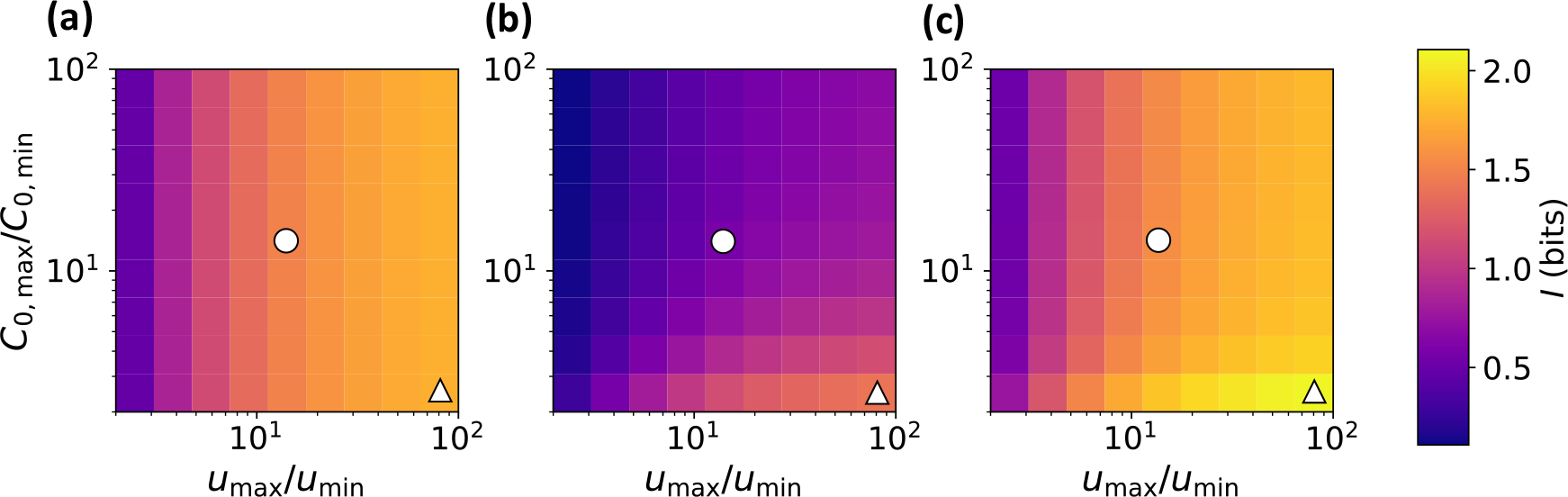
Heat maps of channel capacity for unfolded protein-mediated (a), chaperone-mediated (b) and hybrid (c) mechanisms. The maximal information for each mechanism is computed for different ranges of the uniform prior distribution. The color denotes the channel capacity in bits. In all cases, *C*_0,min_ = 1000 and *u*_min_ = 1000. The upper limits of the input distributions are then varied from 2–100 times the minimum values. In all cases, the unfolded protein-mediated mechanism provides a more informative measurement of unfolded protein concentration than does the chaperone-mediated mechanism. This result is qualitatively insensitive to the minimum values of *C*_0_ and *u*, as well as the number of sensors, *N*_*I*_ (see Supplementary Information). The white circles and triangle mark the input parameters for which the response probability distributions are shown explicitly in Fig. 3.

To better understand the difference in channel capacity between the unfolded protein-mediated and chaperone-mediated sensing mechanisms, we consider the optimized transfer functions (i.e., conditional probability distributions) for a specific prior distribution (Fig. 3). By comparing the conditional probability of activation for the unfolded protein-mediated mechanism (Fig. 3a) with that of the chaperone-mediated mechanism (Fig. 3b), we find that the chaperone-mediated mechanism provides a much more broadly distributed response for a given stimulus. Although the mean responses of the mechanisms are approximately the same, the chaperone-mediated mechanism lacks the specificity provided by the unfolded protein-mediated mechanism, making it far less informative about the number of unfolded proteins in the ER. In particular, the conditional probability of activation for the chaperone-mediated mechanism is skewed toward higher levels of activity. In the following section, we demonstrate that this is due to interference from indirectly measuring the concentration of unfolded protein through the concentration of chaperone.

### Indirect signaling interferes with the measurement of unfolded protein concentration

To better understand why the chaperone-mediated sensing mechanism provides significantly less information about the unfolded protein concentration than the unfolded protein-mediated or hybrid mechanisms, we consider the low-noise limit of the sensor signal transduction. We begin by approximating the transfer function *p*(*n*_*A*_|*u*, *c*) as Gaussian with mean, *μ*(*u*, *C*_0_), equal to the mean of the exact transfer function given in equation (9):

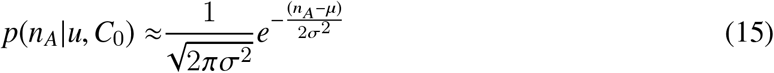

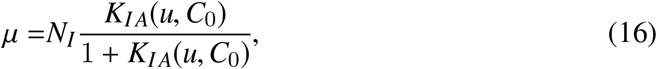

where *σ* is the standard deviation of the Gaussian approximation. Next, we derive an analytical expression for the conditional probability *p*(*n*_*A*_|*u*) for the hybrid mechanism in the limit *σ* → 0.

**Figure 3:**
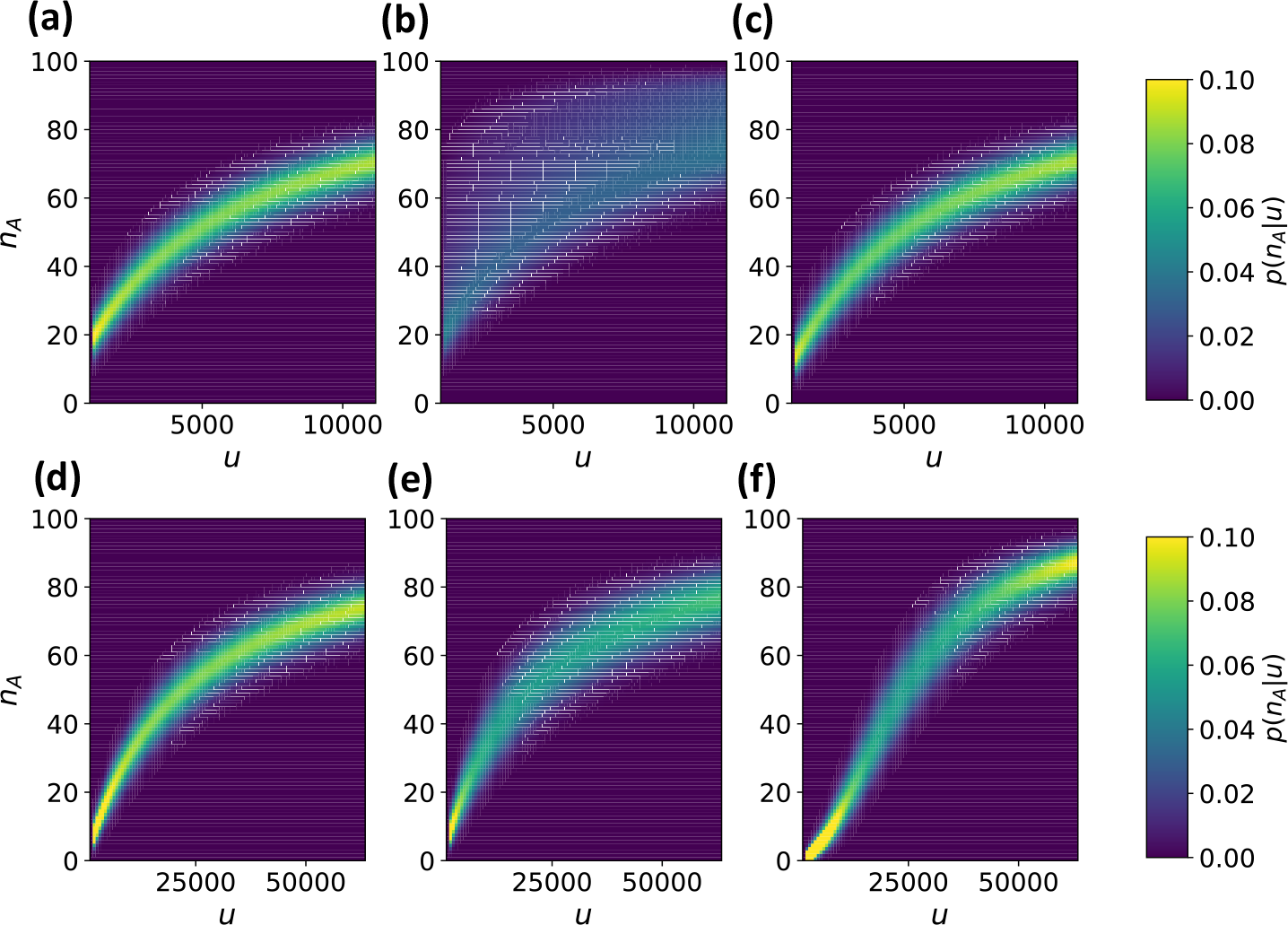
Conditional probabilities of activation as a function of free unfolded protein for the unfolded protein-mediated (a and d), chaperone-mediated (b and e) and hybrid (c and f) mechanisms. The unfolded protein-mediated and hybrid mechanisms provide well defined levels of activation for a given amount of unfolded protein, while the activation of the chaperone-mediated mechanism is much more dispersed. This dispersion greatly limits the information transmission of the chaperone-mediated mechanism. Prior distributions for the top row of panels correspond to those marked by white circles in Fig. 2, and the bottom row of panels correspond to the prior distributions marked by white triangles.

The conditional probabilities of activation for the unfolded protein- and chaperone-mediated models are then found by taking the limits as *β* → 0 and *α* → 0, respectively.

In general, the conditional probability of activation explicitly depends on the amount of chaperone present in the ER and is then given by

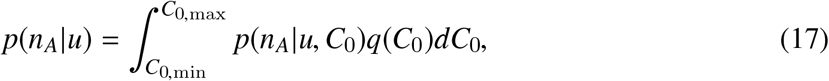

where *q*(*C*_0_) is given by marginalizing the prior distribution over *u*. In the case of the uniform priors used here, this simply results in a uniform distribution between *C*_0,min_ and *C*_0,max_. To approximate the integral on the right-hand-side of equation (17) we employ a saddlepoint approximation (32) (see Supplementary Material for details), valid for small *σ*. This results in an explicit formula for the low-noise approximation to the conditional activation probability:

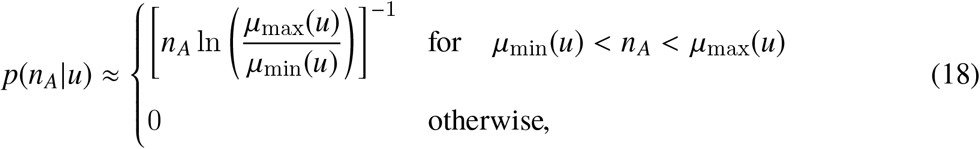

where *μ*_min_(*u*) = *μ*(*u*, *C*_0,max_) and *μ*_max_(*u*) = *μ*(*u*, *C*_0,min_). Hence the width of the the conditional probability distribution depends directly on the range of chaperone concentrations, (*C*_0,min_, *C*_0,max_). In the limit as *μ*_max_ → *μ*_min_, which corresponds to the unfolded protein-mediated mechanism (i.e. *β* → 0), equation (18) reduces to a Dirac delta function centered at the mean value, i.e. *δ*(*n*_*A*_ − *μ*(*u*)). In this case there is no uncertainty about the unfolded protein concentration given a reading of the sensor in the zero-noise limit. However, for both the chaperone-mediated and hybrid mechanism, the dependence of activation on *c* leads to uncertainty even as *σ* → 0. Fig. 4 shows the conditional activation probability distributions for each mechanism mechanism as a function of *u* in the zero-noise limit. For the chaperone-mediated and hybrid mechanisms a degree of uncertainty persists even when the sensor makes a theoretically noise-free measurement due to multiple (*u*, *C*_0_) pairs producing the same sensor output. From equation (18) it is clear that this uncertainty is related to the range of values for *C*_0_ in the prior distribution. When the range set by Δ*C*_0_ shrinks, the precision of the measurement increases, as can be seen by comparing vertically-aligned points in Fig. 2b and c. Hence, indirectly measuring *u* through a mechanism that involves chaperone titration introduces a source of noise that is independent of the stochastic nature of the protein-protein interactions that activate the sensor.

**Figure 4:**
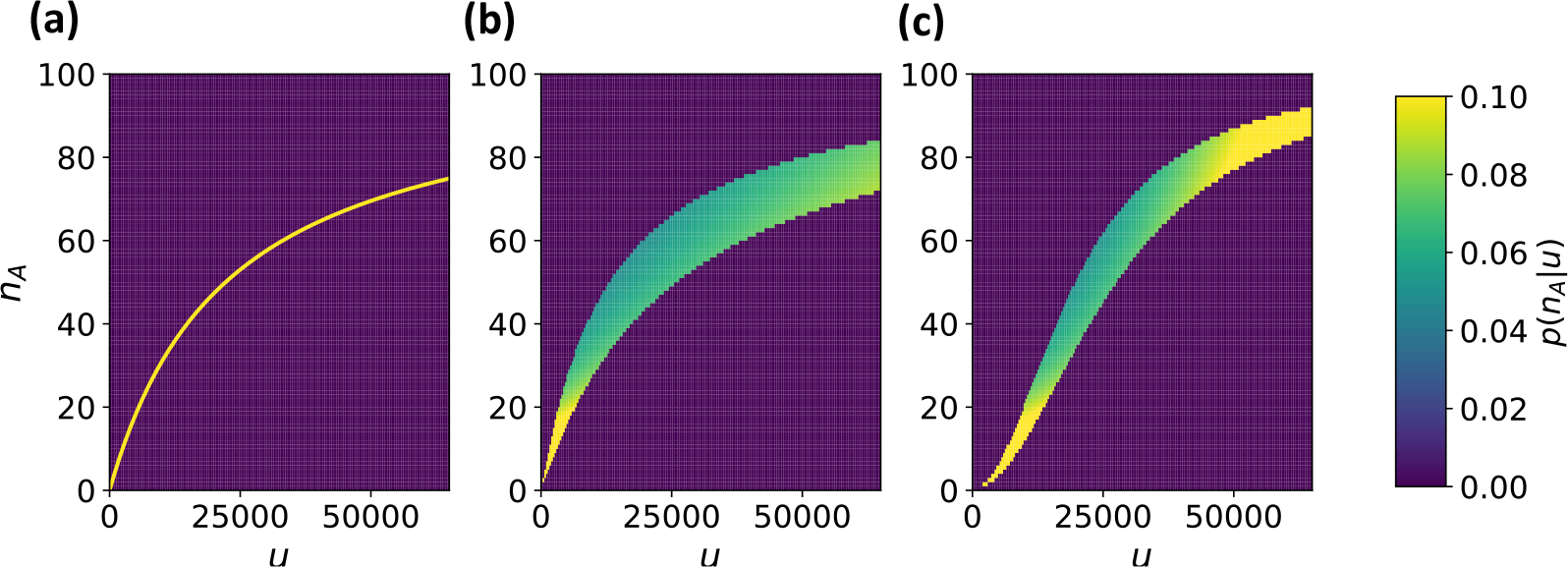
Zero-noise limit of conditional probabilities of activation as a function of free unfolded protein for the unfolded protein-mediated mechanism (a), chaperone-mediated mechanism (b) and the hybrid mechanism (c). Prior distributions and parameters correspond to those marked by white triangles in Fig. 2.

### A hybrid sensing mechanism enhances information transmission by “stretching” the dose-response curve at the expense of increased noise

The information capacity of the hybrid model, shown in Fig. 2c, is always greater than that of the unfolded protein-mediated mechanism. Since the unfolded protein-mediated mechanism is a special case of the hybrid sensing mechanism in which *β* = 0, the hybrid mechanism will always provide at least as much information, but it is not guaranteed that it should outperform the unfolded protein-mediated mechanism. In particular, it is not clear how introducing dependence on an additional random variable, *C*_0_, should increase the sensors ability to measure *u*. The low-noise approximation showed that introducing *C*_0_ into the sensor activation function necessarily obscured the measurement of *u* since multiple values of *u* were then able to produce the same expected output of the sensor. One might expect this to imply that the direct measurement of *u* is the most effective way of measuring *u*.

However, the hybrid mechanism improves the channel capacity beyond the maximal value for the unfolded protein-mediated mechanism. The asymptotic activation probability shown in Fig. 4c offers insight into how this occurs. The hybrid mechanism stretches the range of the sensor compared to the unfolded protein-mediated mechanism, but at the cost of increasing the noise. This is further evidenced by the numerically-calculated mean values and standard deviations of the optimal activation functions for each mechanism, shown in Fig. 5a and b, respectively. The balance of these two competing effects determines the optimal parameterization of the hybrid sensor.

The zero-noise limit sheds additional light on the tradeoff between the range of the sensor and the interference due to measuring stress indirectly. Fig. 5c shows the projection of the surfaces of mean activation onto the *u*−*n*_*A*_ plane. These projections correspond to the same regions for which equation (18) is non-zero. The unfolded protein-mediated mechanism projects onto a single line in the *u*–*n*_*A*_ plane since there is a one-to-one correspondence between *u* and mean sensor activation for this mechanism. The chaperone-mediated and hybrid mechanisms on the other hand lack this one-to-one correspondence and a single mean output value corresponds to a range of values of *u*. The hybrid mechanism, however, effectively overcomes this added uncertainty by increasing the range of mean activation. In summary, the hybrid mechanism increases uncertainty due to indirectly measuring *u* through the chaperone concentration in order to extend the sensor operating range. This tradeoff allows the hybrid mechanism to transmit maximal information about *u*.

### A hybrid sensing mechanism also increases information about free chaperone concentration

Thus far, we have considered *u* to be the measure of stress that the cell aims to monitor. However, this need not be the case. For example, an ER with very few unbound chaperones indicates that the folding capacity is nearly exceeded by client unfolded protein demand and action must be taken to maintain proteostasis. Furthermore, it has been shown that a UPR that responds to the concentration of free chaperone, as opposed to free unfolded protein, can provide a more efficient response to acute stress (25). In this section, we demonstrate that the hybrid mechanism can provide a more precise reading of free chaperone concentration in the same way it was able to provide a more precise measurement of unfolded protein concentration.

**Figure 5:**
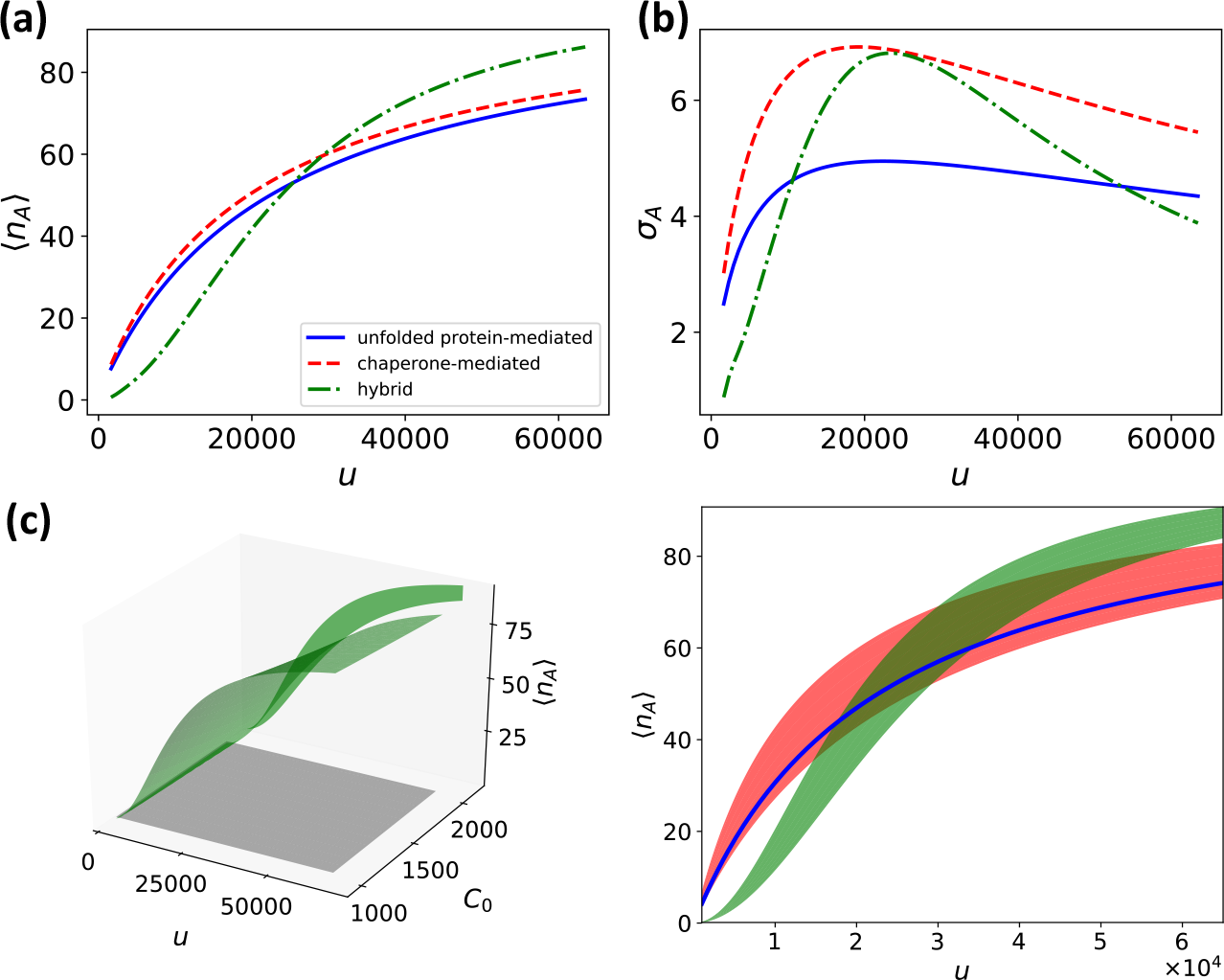
Panels (a) and (b) show the mean activation and standard deviation for each mechanism as a function of unfolded protein. Panel (c) shows the surface of mean activation for the hybrid mechanism (green surface, left panel), over the uniform prior distribution (grey square), along with the projection of the hybrid mean activation surface onto the *u*−*n*_*A*_ plane. On the right, the projection of surfaces of mean activation onto the *u*−*n*_*A*_ plane are plotted for each mechanism. The hybrid mechanism (green shaded region), which provides the most informative signal, increases the range of activation at the expense of greater noise compared to the unfolded protein-mediated sensor (blue line). The chaperone-mediated mechanism (red shaded region) projects onto a broad area of the input-output space, making it a poor sensor of unfolded protein. Parameters and input distributions correspond to those marked by the white triangles in Fig. 2.

Fig. 6 shows optimized conditional activation probabilities for each mechanism as a function of free chaperone. When the aim of the sensor is to measure free chaperone concentration, the unfolded protein-mediated sensor suffers from the same interference effect that the chaperone-mediated sensor suffered when the quantity of interest was unfolded protein. Both the chaperone-mediated sensor and the hybrid sensor provide relatively reliable measurements of the chaperone concentration. Again, it is possible to construct a low-noise approximation for the conditional activation probability (see Supplementary Material):

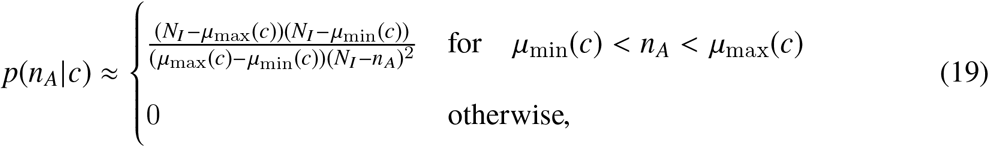

where now *μ*_min_(*c*) = *μ*(*c*, *U*_0,min_) and *μ*_max_(*c*) = *μ*(*c*, *U*_0,max_).

Equation (19), evaluated at optimized parameters for each mechanism, is shown in Fig. 6 (bottom row). Analogously to the case where *u* was the quantity of interest, the hybrid mechanism is able to provide more information about the concentration of free chaperone than the chaperone-sensing mechanism by stretching the range of activation of the sensor for the same input range of *c*. As shown in Fig. 7a and b, this is again the result of allowing some additional uncertainty in the output for specific inputs in exchange for a greater range of outputs. Fig. 7c provides a geometrical interpretation: identical values of *c* can produce different outputs, *n*_*A*_, for the unfolded protein-mediated and hybrid mechanisms. Geometrically, this corresponds to the surface of mean activation projecting onto a region in the *c*−*n*_*A*_ plane with a finite area. Mitigating this uncertainty by increasing the range of the sensors response, the hybrid sensing mechanism allows for more informative measurements of the free chaperone concentration than a mechanism that only responds directly to free chaperones. This result, together with the similar result for measuring unfolded protein concentration, demonstrates that directly measuring the quantity of interest does not necessarily provide the most information about that quantity.

**Figure 6:**
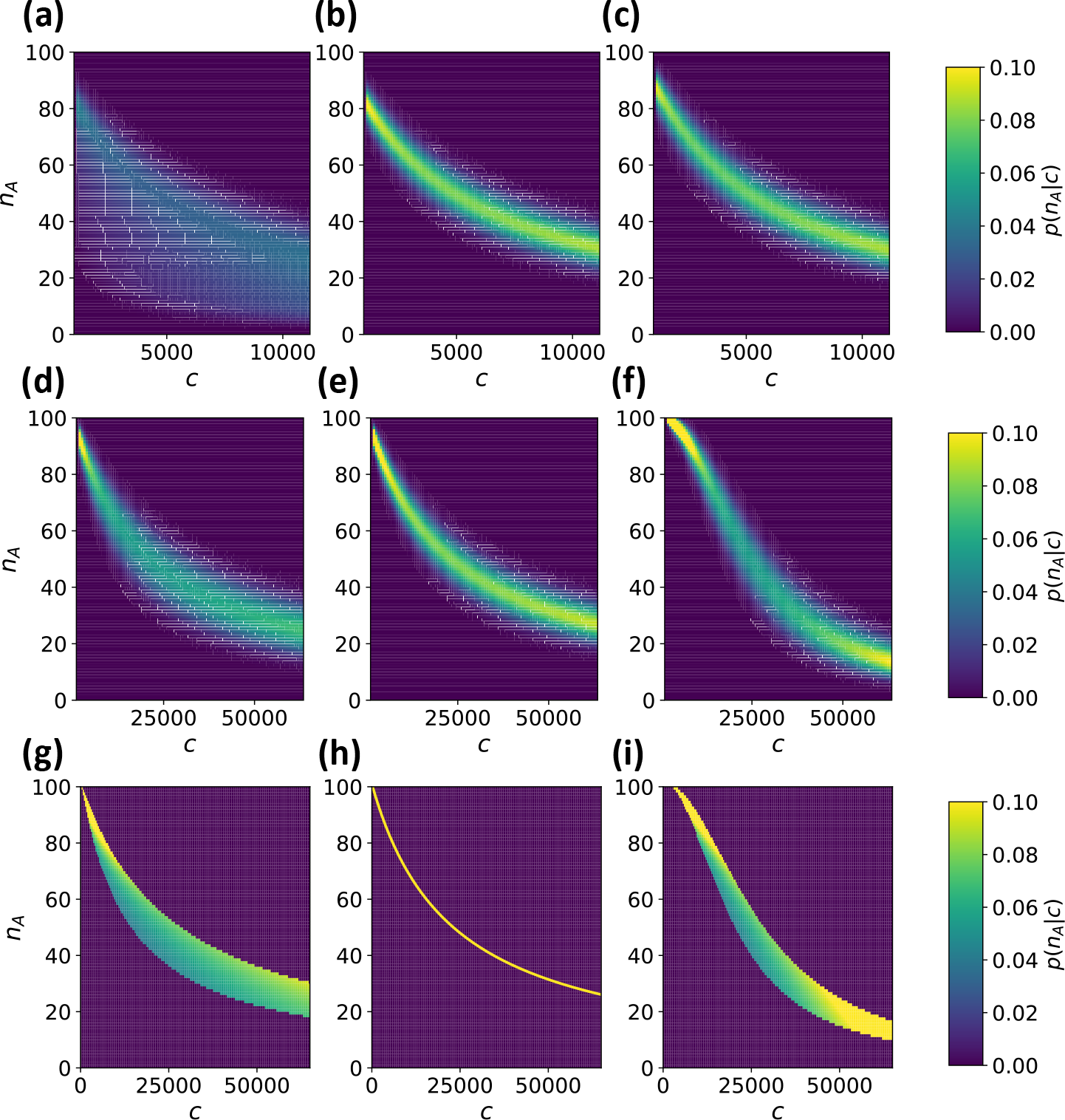
Conditional probabilities of activation as a function of free chaperone for the unfolded protein-mediated (left column), chaperone-mediated (middle column) and hybrid (right column) mechanisms. The top two rows show numerically calculated conditional probabilities of optimized parameter sets for two different prior distributions. The bottom row shows the low-noise approximation given in equation (19) for the same prior and parameters used in the middle row. The chaperone-mediated and hybrid mechanisms provide well defined levels of activation for a given amount of free chaperone, while the activation of the unfolded protein-mediated mechanism is much more dispersed. This dispersion greatly limits the information transmission of the unfolded protein-mediated mechanism. The plotted results are for uniform prior distributions with ranges *U*_0,min_ = 10^3^, *U*_0,max_ = 11.4 × 10^3^, *c*_min_ = 10^3^ and *c*_max_ = 11.4 × 10^3^ for the top row, and *U*_0,min_ = 10^3^, *U*_0,max_ = 2 × 10^3^, *c*_min_ = 10^3^ and *c*_max_ = 64.7 × 10^3^ for the bottom two rows. The mutual information for priors with other ranges are provided in the Supplementary Material.

**Figure 7:**
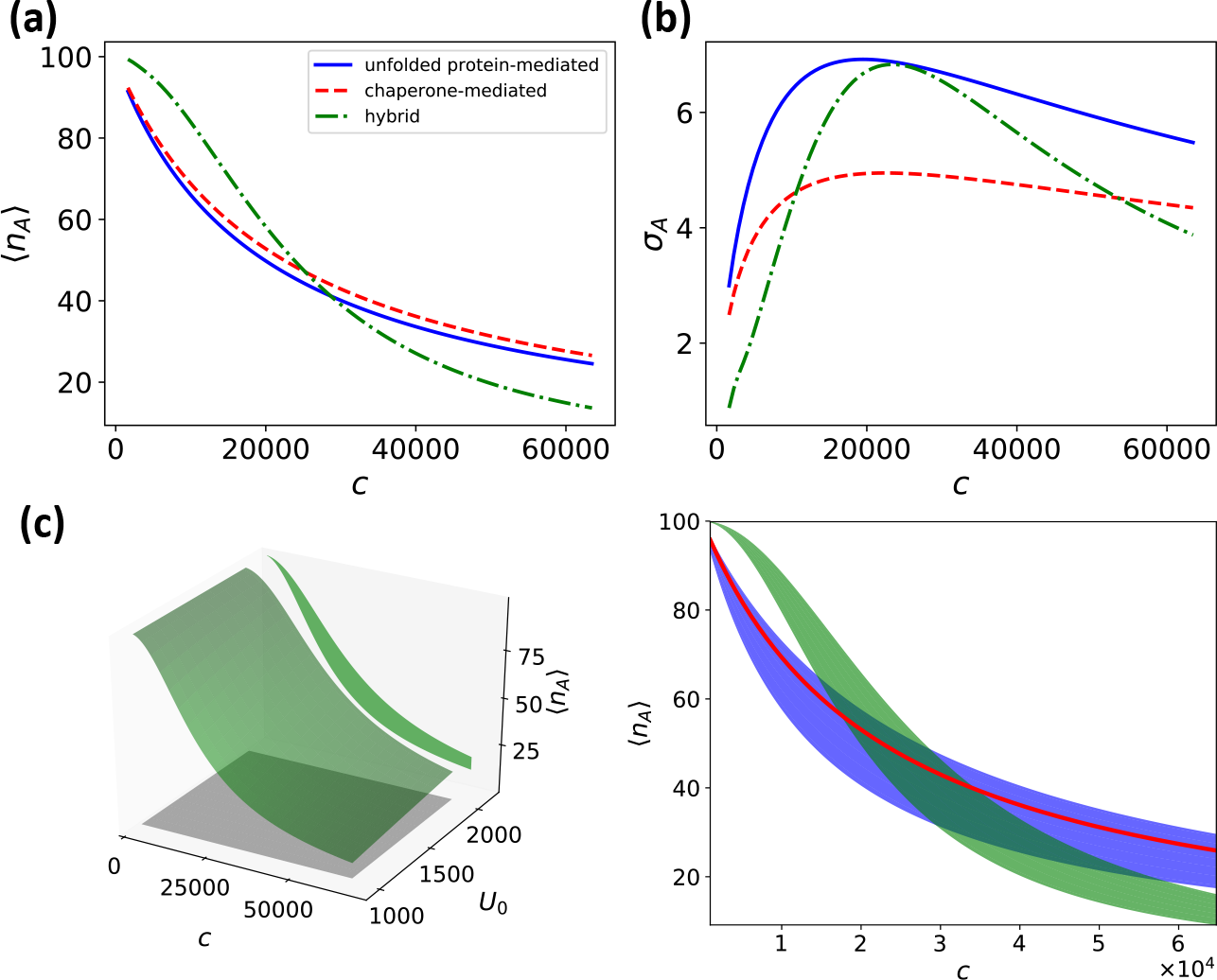
Panels (a) and (b) show the mean responses and standard deviations of each mechanism as a function of free chaperone. Panel (c) shows the surface of mean activation for the hybrid mechanism (green surface, left panel), over the uniform prior distribution (grey square), along with the projection of the hybrid mean activation surface onto the *c*−*n*_*A*_ plane. On the right, the projection of surfaces of mean activation onto the *c*−*n*_*A*_ plane are plotted for each mechanism. The hybrid mechanism (green shaded region) increases the range of activation at the expense of greater noise as compared to the chaperone-mediated sensor (red line). The unfolded protein-mediated mechanism (blue shaded region) projects onto a broad area of the input-output space, making it a poor sensor of available chaperone.

## Discussion

ER stress is monitored by stress sensing proteins in the ER membrane that are both activated by unfolded protein ligands and suppressed by unbound chaperones. However, it has not been established why such a mechanism evolved. We hypothesized that this hybrid mechanism of stress sensing could provide more information about the state of the ER than measuring only unfolded proteins or chaperones, while still allowing the UPR to take advantage of the added efficiency and buffering of sensors that respond to depleted chaperone.

Our results indicates that a sensor that is suppressed by chaperone provides less information about unfolded protein concentration in the ER than a sensor that is activated by unfolded proteins. In particular, this is because multiple input pairs (*u*, *C*_0_) can produce the same output of the chaperone-mediated sensor even when the signal is free of noise. This introduces inherent ambiguity into the sensor output with regards to the concentration of free unfolded proteins within the ER, i.e. ER stress. A mechanism that combines both direct unfolded protein sensing and chaperone sequestration of sensors is capable of providing more information than the unfolded protein-mediated mechanism alone. To do so, the hybrid sensor allows for a small increase in ambiguity of the mean sensor output for a given concentration of unfolded protein in order to extend the range of outputs.

Analogously, if the sensor evolved to measure the concentration of free chaperone in the ER, which is an alternative measure of ER stress, the chaperone-mediated sensor provides substantially more information than the unfolded-protein sensor. The hybrid sensor then further increases the channel capacity of the chaperone-mediated sensor. Previous studies (24; 25) have shown that the chaperone-mediated sensing mechanism provides a benefit in terms of the efficiency of the UPR when responding to acute stress events. However, the precision with which the sensor determines the level of stress in the ER was not considered. The present analysis illustrates that combining chaperone-mediated suppression with unfolded-protein activation can increase the amount of information about ER stress transmitted out of the ER lumen.

By providing a more informative measurement of the stress level within the ER, the hybrid mechanism would allow for a more finely-tuned UPR. Precise control over the UPR is important to organism fitness due to the substantial metabolic cost of maintaining proteostasis. For example, the chaperone BiP is present at one of the highest copy numbers of any protein in eukaryotic cells, ≈ 3 × 10^5^ in yeast (33) and ≈ 2 × 10^7^ in unstressed HeLa cells (34). Upon stress, the copy number can increase more than 10 fold (34). Decreasing the amount of chaperone required to mitigate an acute stress event can therefore make a substantial contribution to cellular energy expenditure. This is particularly true for secretory cells, such as pancreatic *β*-cells, where proinsulin mRNA translation rates reach approximately 10^6^ molecules per minute in response to glucose stimulation (35; 36; 37), all of which must be processed in the ER. Overall, up to 20% of known genes go through transcriptional changes when the UPR is activated (38). Additionally, for large stresses, the UPR induces apoptosis, further increasing the importance of reliable and precise ER stress measurements. Hence, the importance of precise UPR control provides a rationalization of the ER stress sensing mechanism. A sensor that is repressed by available chaperones provides a relatively simple mechanism for monitoring and responding to chaperone abundance in the ER, and is able to capitalize on the advantages of using free chaperone as a measure of stress. However, the precision of this mechanism can then be further enhanced by incorporating a direct interaction with unfolded proteins into the activation mechanism.

This reasoning depends on the implicit assumption that the information capacity of the stress sensing network and the metabolic efficiency with which the network responds to stress are the main drivers of stress sensor evolution. While we acknowledge this to be an assumption, we believe it to be valid for two reasons. First, the metabolic cost of mitigating protein stress and maintaining protein homeostasis are substantial, especially for secretory cells. Hence, a more frugal use of chaperone upregulation in response to stress can significantly reduce cellular energy expenditure. Second, for the cell to take advantage of the more efficient stress response, the signal must be informative regarding the level of stress. In particular, it is essential for the first step in the signal transduction pathway – the measurement of stress in the ER – to transmit maximal information since the information processing inequality ensures that information will only be further degraded as it passes along the signaling pathway.

While incorporating the direct activation of the sensor by unfolded proteins enhances the information about ER stress beyond the capacity of the chaperone-mediated sensor alone, this may be only one of several benefits provided by direct unfolded protein-sensor interactions. For example, the binding of the unfolded protein could stabilize clusters of signaling molecules into long-lived signaling foci, changing the dynamics of the response (24). Analysis of the binding groove formed by Ire1 dimers suggests that the peptide sequences that are recognized by the signaling complex are only partially overlapping with those which act as substrates for the chaperone BiP (19). This could indicate that direct protein binding provides supplementary information about unfolded proteins in the ER that is not incorporated in the concentration of available BiP. However, regardless of whether these effects are also important, our analysis reveals that incorporating the unfolded protein directly into the signaling mechanism can enhance the measurement of the free chaperone in the ER.

The additional information capacity of the hybrid sensing mechanism represents a specific instance of a more general feature of signal integration in biochemical networks. Namely, it is possible to increase the precision with which one component of a multi-component system can be sensed by incorporating the signal from another coupled component. This will be the case for any instance in which the aim is to monitor the components of a bimolecular reaction, including enzyme catalyzed reactions. Given the prevalence of such reactions in cellular biochemistry, we expect the results presented here to extend beyond the case of ER stress sensing.

## Conclusion

This work provides two main results regarding stress sensing in the ER. First, on its own, the chaperone-mediated sensing mechanism provides a poor estimate of the concentration of unfolded proteins in the ER, but a rather precise measurement of the concentration of available chaperone. Hence, it appears that cells might have evolved to respond primarily to the depletion of available chaperone as opposed to unfolded protein copy number. Second, integrating the signal from free chaperones with direct sensor activation by unfolded proteins can improve the information about chaperone availability within the ER. Together, these results further our understanding of how cells monitor and maintain protein folding homeostasis within the ER.

## Supporting information

Supplemental Material to Numerical Simulations

## Authors’ Contributions

WS and SS conceived the study. WS generated numerical results. WS and JE analyzed the model. WS, JE, and SS interpreted the results and wrote the manuscript. All authors gave approval of the final version of the manuscript.

## Data Accessibility

All codes used to generate and analyze results are available at https://github.com/santiago-schnell/Information-Processing-ER-Stress-Sensors

