## Supplemental Material to Numerical Simulations for "Information processing by endoplasmic reticulum stress sensors"

This Supplementary Material consists of three sections. Section 1 details the numerical calculation of the mutual information and the procedure used to determine parameters that maximize the mutual information. Section 2 provides a detailed derivation of the low-noise approximations for the conditional activation probability distributions. Lastly, Section 3 presents numerical calculations showing that the results presented in the manuscript are robust to parameter choice.

#### 1 Numerical calculation of mutual information

##### Optimization of mutual information

Maximization of the mutual information for each mechanism is performed using the Limited-memory Broyden-Fletcher-Goldfarb-Shanno for Bounded (L-BFGS-B) constrained optimization algorithm [1], as implemented in SciPy v1.1.0. Additionally, we augment this algorithm with a multi-start procedure as follows. For each optimization parameter, we assign three initial values of $10^{-4}$, 1, and $10^{4}$ and maximize the mutual information for each possible combination of initial conditions. Following this procedure, we also start the optimization of the hybrid mechanism at each of the parameter sets that maximize the information of the two special cases. For each mechanism, the parameterization that maximizes the information over all initial conditions is taken to be the optimal parameterization. This procedure allows for broad sampling of the parameters space and results in maxima that are robust to specific choices of initial conditions for the optimization procedure.

##### Convergence of simulations upon discretization refinement

The numerical calculation of the mutual information requires the discretization of the prior probability distributions. The two-dimensional uniform priors considered in this work are discretized into an $N_X \times N_Y$ rectangular array of equally spaced bins, where $(X, Y) = (u, C_0)$ for the case in which $u$ is the quantity of interest, and $(X, Y) = (c, C_0)$ when $c$ is the quantity of interest. To compute the mutual information the initial parameterization of the priors are transformed into distributions over $(u, c)$. This requires rebinning of the discrete priors, i.e., transforming $(u, C_0) \to (u, c)$ or $(c, U_0) \to (u, c)$. The number of bins used for the rebinning is $N_B$. In all cases, we choose $N_X = N_Y = N_B = N$. To ensure that the discretization is not introducing significant error into the calculation of the mutual information we have tested the convergence of the mutual information as a function of increasing $N$. In particular, we calculate the error as

$$\text{Error}(N) = \frac{|MI(N) - MI(N_{\max})|}{MI(N_{\max})}, \tag{S1}$$

where $MI$ is the mutual information and $N_{\max}$ is the finest discretization tested, which is $N = 200$. Fig. S1 shows that the error becomes smaller than 0.5% for all mechanisms in the case of $u$ being the quantity of interest when $N = 100$. The same is shown for the case in which $c$ is the quantity of interest in Fig. S2. Hence we have used $N = 100$ for the discretization throughout this work.

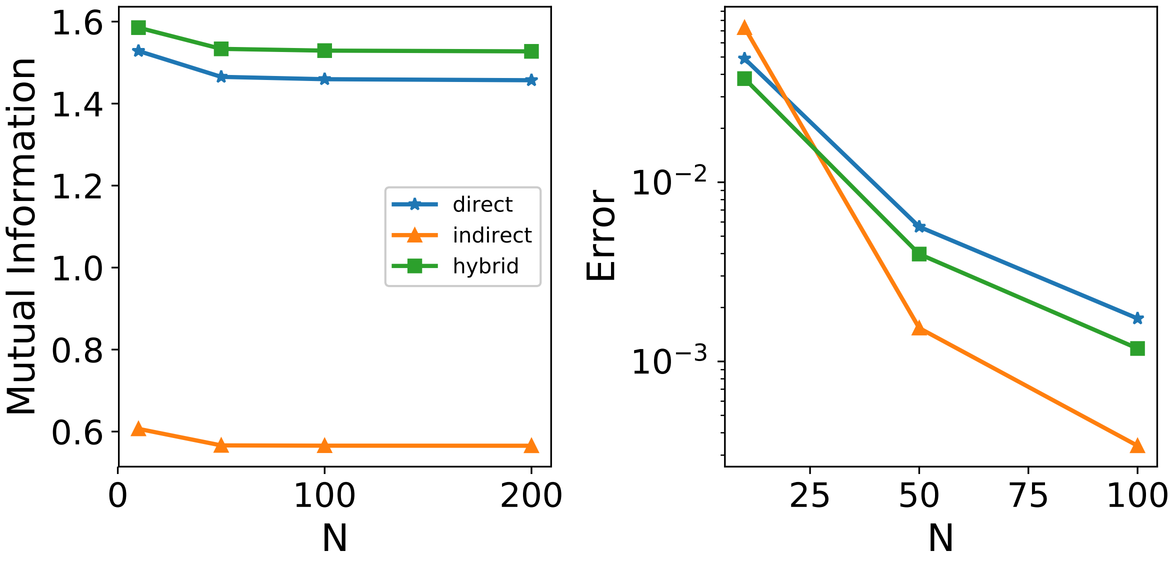


Supplementary Figure S1: Mutual information (left) and error in mutual information (right) as a function of number of bins, $N$, used to discretize the prior distribution for the case in which free unfolded protein, $u$, is the quantity of interest. For $N = 100$, which is the quantity used for all results in the main text, the percent error is below 0.5%. The data points are calculated using the following parameters: $u_{\min} = 10^3$, $u_{\max} = 10^4$, $C_{0,\min} = 10^3$, $C_{0,\max} = 10^4$, $N_I = 100$.

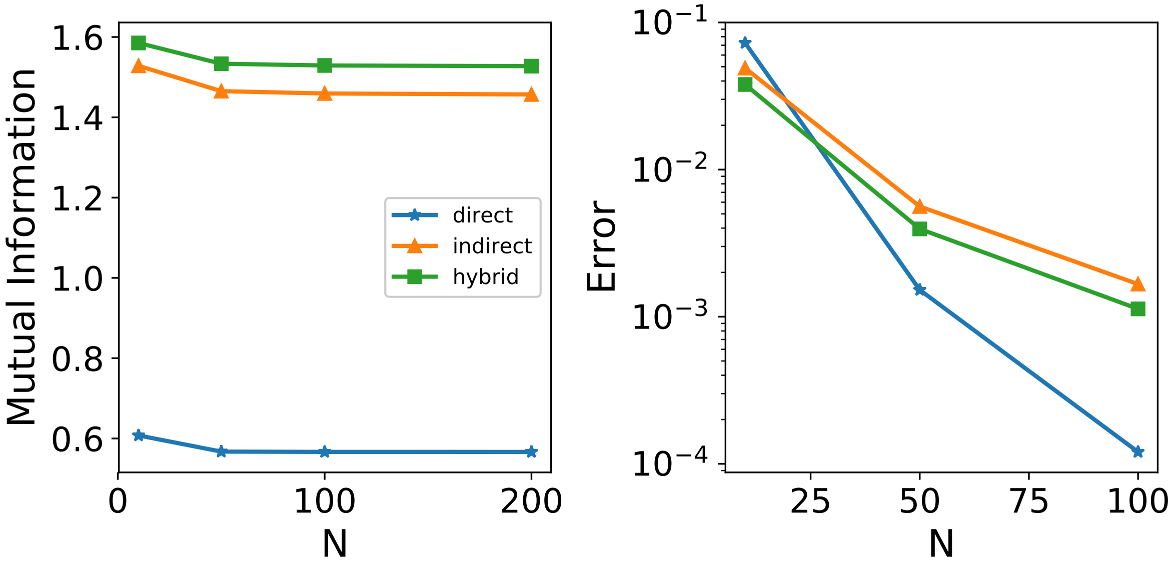


Supplementary Figure S2: Mutual information (left) and error in mutual information (right) as a function of number of bins, $N$, used to discretize the prior distribution for the case in which free chaperone, $c$, is the quantity of interest. For $N = 100$, which is the quantity used for all results in the main text, the percent error is below 0.5%. The data points are calculated using the following parameters: $c_{\min} = 10^3$, $c_{\max} = 10^4$, $C_{0,\min} = 10^3$, $C_{0,\max} = 10^4$, $N_I = 100$.

#### 2 Asymptotic approximation for the conditional activation probability

This section provides a detailed derivation of the low-noise approximation for the conditional activation probability distributions given by Equation (18) (case 1) and Equation (19) (case 2) in the main text.

##### Case 1: Quantity of interest is $u$

To approximate the integral on the right-hand-side of Equation (17) we will use a saddlepoint approximation, but first it is useful to change coordinates from $(u, C_0) \to (u, \mu)$ [2]. Equation (17) then becomes

$$p(n_A|u) = \int_{\mu_{\min}}^{\mu_{\max}} p(n_A|u,\mu) q(\mu|u) \left(\frac{\partial \mu}{\partial C_0}\right)^{-1} d\mu, \tag{S2}$$

where $\mu_{min}(u) = \mu(u, C_{0,\max})$ and $\mu_{max}(u) = \mu(u, C_{0,\min})$, and the explicit dependence of $\mu$ on $u$ and $C_0$ has been omitted. $q(\mu|u)$ is the prior distribution transformed into $(u, \mu)$ coordinates and conditioned on $u$ given by

$$q(\mu|u) = \begin{cases} \dfrac{1}{\Delta\mu} & \text{for} \quad \mu_{\min} < \mu < \mu_{\max} \\ 0 & \text{otherwise} \end{cases}$$

with $\Delta\mu = \mu_{\max} - \mu_{\min}$. The right-hand-side of equation (S2) can be readily recast in a form appropriate for a saddlepoint approximation

$$p(n_A|u) = \int_{-\infty}^{\infty} \frac{f(\mu|u)}{\sqrt{2\pi\sigma^2}} e^{-\frac{1}{\sigma^2}g(\mu)}, \tag{S3}$$

where

$$f(\mu|u) = -q(\mu|u) \left(\frac{\partial\mu}{\partial C_0}\right)^{-1}, \tag{S4}$$

$$g(\mu) = \frac{(n_A - \mu)^2}{2}, \tag{S5}$$

and the bounds of the integral (after switching their order) can be extended to $(-\infty, \infty)$ since the prior $p(\mu)$ is zero everywhere outside of the support. The saddlepoint approximation relies on a local maxima of the integrand, which is given by solving $\partial g/\partial\mu|_{\mu=\mu_c} = 0$, leading to $\mu_c = n_A$. In the limit $\sigma \to 0$, The saddlepoint approximation of Equation (S3) is then

$$\begin{aligned} p(n_A|u) &\approx e^{-\frac{1}{\sigma^2}g(\mu_c)} \frac{f(\mu_c|u)}{\sqrt{2\pi\sigma^2}} \sqrt{\frac{2\pi\sigma^2}{(\partial^2 g/\partial\mu^2)|_{\mu_c}}} \\ &= f(n_A|u). \end{aligned} \tag{S6}$$

Hence, all that is needed now is to compute the explicit form of $f(\mu|u)$. For the hybrid mechanism, $\mu$ is given explicitly by

$$\mu(u, C_0) = \frac{N_I A(u)}{1 + \beta C_0 + A(u)}, \tag{S7}$$

where

$$A(u) = \gamma \left[\alpha u^2 + (1+\alpha)u + 1\right]. \tag{S8}$$

Computing the Jacobian for the coordinate transformation, we obtain

$$\frac{\partial\mu}{\partial C_0} = -\frac{N_I \beta A(u)}{(1 + \beta C_0 + A(u))^2}. \tag{S9}$$

Noting that

$$C_0(\mu, u) = \frac{1}{\beta}\left(\frac{N_I A(u)}{\mu} - (1 + A(u))\right), \tag{S10}$$

we obtain

$$\left(\frac{\partial\mu}{\partial C_0}\right)^{-1} = -\frac{N_I A(u)}{\beta\mu}. \tag{S11}$$

Then, substituting Equation (S11) and the uniform prior distribution into Equation (S4), we obtain

$$f(\mu|u) = \begin{cases} \dfrac{N_I A(u)}{\Delta\mu\beta\mu} & \text{for} \quad \mu_{\min} < \mu < \mu_{\max} \\ 0 & \text{otherwise} \end{cases}$$

where $\mu_{min}(u) = \mu(u, C_{0,\max})$ and $\mu_{max}(u) = \mu(u, C_{0,\min})$. The final step is to normalize $f(\mu|u)$ to make it a probability density function. The normalization constant is

$$Z(u) = \int_{\mu_{min}}^{\mu_{max}} f(\mu|u) d\mu \tag{S12}$$

$$= \frac{N_I A(u)}{\Delta\mu\beta} \left[\ln \mu_{max}(u) - \ln \mu_{\min}(u)\right]. \tag{S13}$$

Hence the low-noise approximation to the conditional activation probability for the hybrid mechanism is

$$\begin{aligned} p(n_A|u) &\approx \frac{f(n_A|u)}{Z(u)} \\ &= \begin{cases} \left[n_A \ln\left(\dfrac{\mu_{max}(u)}{\mu_{\min}(u)}\right)\right]^{-1} & \text{for} \quad \mu_{min}(u) < n_A < \mu_{max}(u) \\ 0 & \text{otherwise,} \end{cases} \end{aligned} \tag{S14}$$

as given by Equation (18) in the main text.

#### Case 2: Quantity of interest is $c$

For the case in which $c$ is the quantity of interest, the procedure remains essentially the same, though some expressions differ. In particular, the saddle point approximation leads to

$$p(n_A|c) \approx \frac{f_c(n_A|c)}{Z_c(c)} \tag{S15}$$

$$= \frac{1}{Z_c(c)} \left[ -q(\mu|c) \left(\frac{\partial \mu}{\partial U_0}\right)^{-1} \right]\Bigg|_{\mu=n_A}, \tag{S16}$$

where $Z_c$ is a normalization constant. $\mu$ can be written in terms of $(c, U_0)$ as

$$\mu(c, U_0) = \frac{N_I \gamma \left(\alpha U_0 + 1 + c\right)}{1 + c\left(1 + \beta\right) + \beta c^2 + \gamma\left(\alpha U_0 + 1 + c\right)}. \tag{S17}$$

Differentiating Equation (S17) with respect to $U_0$ and transforming coordinates in order to eliminate $U_0$ leads to

$$\left(\frac{\partial \mu}{\partial U_0}\right)^{-1} = \frac{N_I(1+c)(1+\beta c)}{\gamma\alpha(N_I - \mu)^2}. \tag{S18}$$

Combining Equation (S18) with the uniform prior distribution give

$$f_c(\mu|c) = \begin{cases} \dfrac{N_I(1+c)(1+\beta c)}{\Delta\mu\gamma\alpha(N_I-\mu)^2} & \text{for} \quad \mu_{\min} < \mu < \mu_{\max} \\ 0 & \text{otherwise} \end{cases}$$

where now $\mu_{\min} = \mu(c, U_{0,\min})$ and $\mu_{\max} = \mu(c, U_{0,\max})$. The normalization constant is given by

$$\begin{aligned} Z_c(c) &= \frac{N_I(1+c)(1+\beta c)}{\Delta\mu\gamma\alpha} \int_{\mu_{\min}}^{\mu_{\max}} \frac{1}{(N_I-\mu)^2} d\mu \\ &= \frac{N_I(1+c)(1+\beta c)}{\Delta\mu\gamma\alpha} \frac{\Delta\mu}{(N_I-\mu_{\max})(N_I-\mu_{\min})}, \end{aligned} \tag{S19}$$

which leads to

$$\frac{f_c(\mu|c)}{Z_c(c)} = \begin{cases} \dfrac{(N_I-\mu_{\max})(N_I-\mu_{\min})}{\Delta\mu(N_I-\mu)^2} & \text{for} \quad \mu_{\min} < \mu < \mu_{\max} \\ 0 & \text{otherwise.} \end{cases}$$

Evaluating Equation (S20) at $\mu = n_A$ gives the low-noise approximation for the conditional activation probability presented in Equation (19) of the main text.

### 3 Robustness of results to model parameters

#### Effect of sensor copy number on mutual information

In the manuscript, we assumed the sensor copy number to be $N_I = 100$. This is on the order of the Ire1 copy number found in yeast [3]. However, here we demonstrate that the main qualitative results of the work are independent of this choice. Fig. S3 shows heatmaps of the maximal mutual information between $u$ and $n_A$ for each mechanism when $N_I = 50$ (top row) and $N_I = 200$ (bottom row). While the number of sensor molecules increases the amount of information for a given prior distribution, it does not change the observation that the unfolded protein-mediated sensor and hybrid sensor mechanisms provide significantly more information than the chaperone-mediated sensing mechanism.

Similarly, Fig. S4 shows heatmaps of the maximal mutual information between $c$ and $n_A$ for each mechanism when $N_I = 50$ and $N_I = 200$. Again, the observation that the chaperone-mediated sensor and the hybrid mechanism perform significantly better than the unfolded protein-mediated mechanism remains true. Hence, the number of sensors can increase the channel capacity of the sensing system, but does not change the relative performance of the different mechanisms.

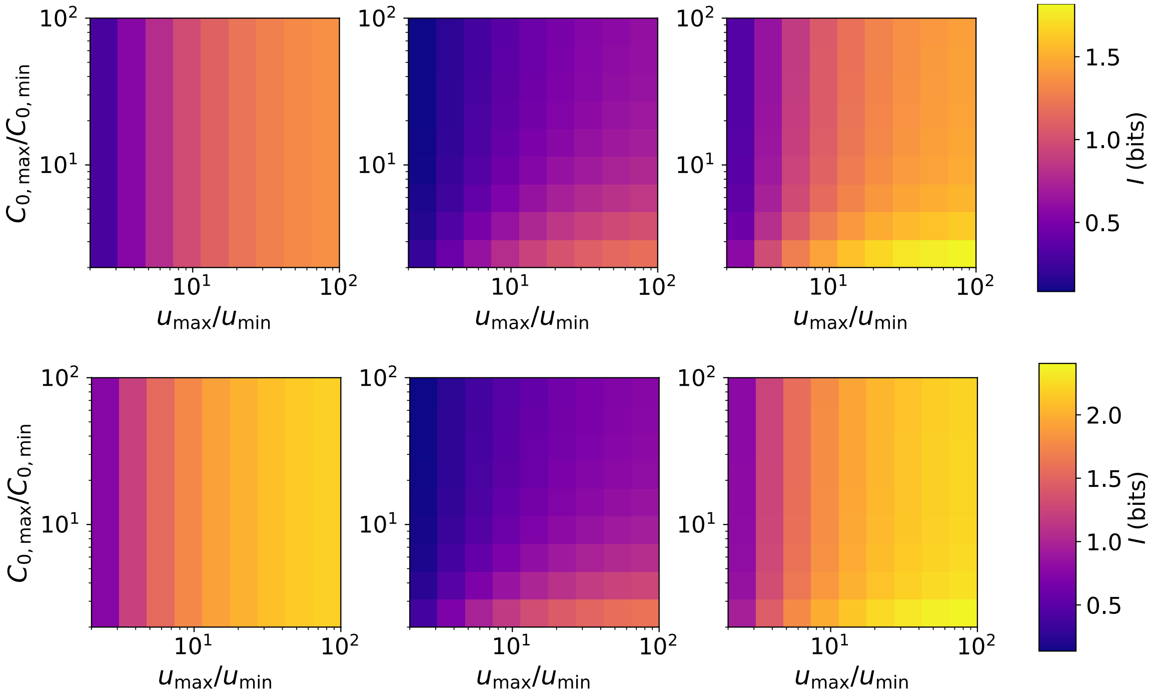


Supplementary Figure S3: Heat maps showing mutual information for different ranges of the uniform prior distribution when $u$ is the quantity of interest. The top row is for $N_I = 50$ and the bottom row is for $N_I = 200$. The left, middle and right columns correspond to the unfolded protein-mediated, chaperon-mediated and hybrid mechanisms, respecively. The data points are calculated using the following parameters: $u_{\mathrm{min}} = 10^3$, $C_{0,\mathrm{min}} = 10^3$.

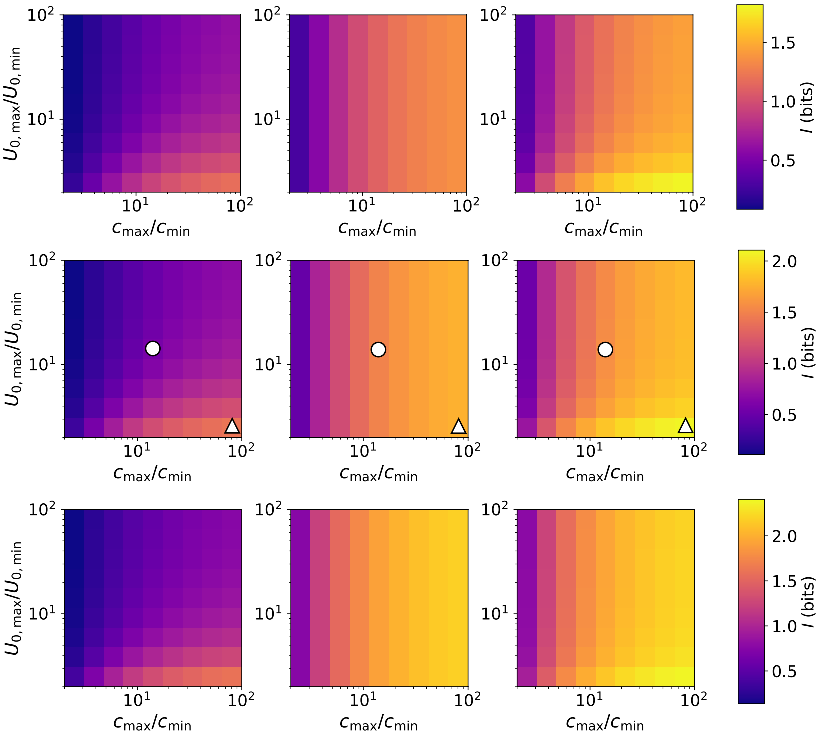


Supplementary Figure S4: Heat maps showing mutual information for different ranges of the uniform prior distribution when $c$ is the quantity of interest. The top row is for $N_I = 50$, middle row is for $N_I = 100$ and the bottom row is for $N_I = 200$. The left, middle and right columns correspond to the unfolded protein-mediated, chaperon-mediated and hybrid mechanisms, respecively. The white cirles and white triangles correspond to the conditional activation probabilities shown in Fig. 5 of the main text. The data points are calculated using the following parameters: $c_{\text{min}} = 10^3$, $U_{0,\text{min}} = 10^3$.

#### Effect of $u_{\text{min}}$, $c_{\text{min}}$ and $C_{0,\text{min}}$ on mutual information

The heatmaps shown in Fig. 1 of the main text assume values for the lower limits of the uniform prior distributions in $(u, C_0)$ and $(c, C_0)$, respectively, and show the effect of varying the upper limits. Here we show that the choice of lower limits $(u_{\text{min}}, C_{0,\text{min}})$ and $(c_{\text{min}}, C_{0,\text{min}})$ does not qualitatively change the results. Fig. S5 shows heat maps for the cases $(u_{\text{min}}, C_{0,\text{min}}) = (10^2, 10^2)$ (top row) and $(u_{\text{min}}, C_{0,\text{min}}) = (10^4, 10^4)$ (bottom row) when $u$ is the quantity of interest. Comparing these data with the heatmap in Fig. 1 in the main text shows that the lower limit on the prior distribution does not alter the relative effectiveness of the sensing mechanisms. In fact, the magnitude of the mutual information is approximately equal for corresponding points on the heat maps, regardless of $(u_{\text{min}}, C_{0,\text{min}})$. This is the result of the parameters dictating the shape of the response curve being optimized for each case independently, effectively normalizing the absolute scale of the prior distribution.

Similarly, Fig. S6 shows the effect of varying $c_{\rm min}$ and $U_{0,\rm min}$ on the channel capacity when $c$ is the quantity of interest. Changing the absolute value of the range of the prior does not alter the result that the chaperone-mediated sensor and the hybrid sensor outperform the unfolded-protein mediated sensor. Hence, the results of this work are independent of the choice of $(c_{\rm min}, U_{0,\rm min})$.

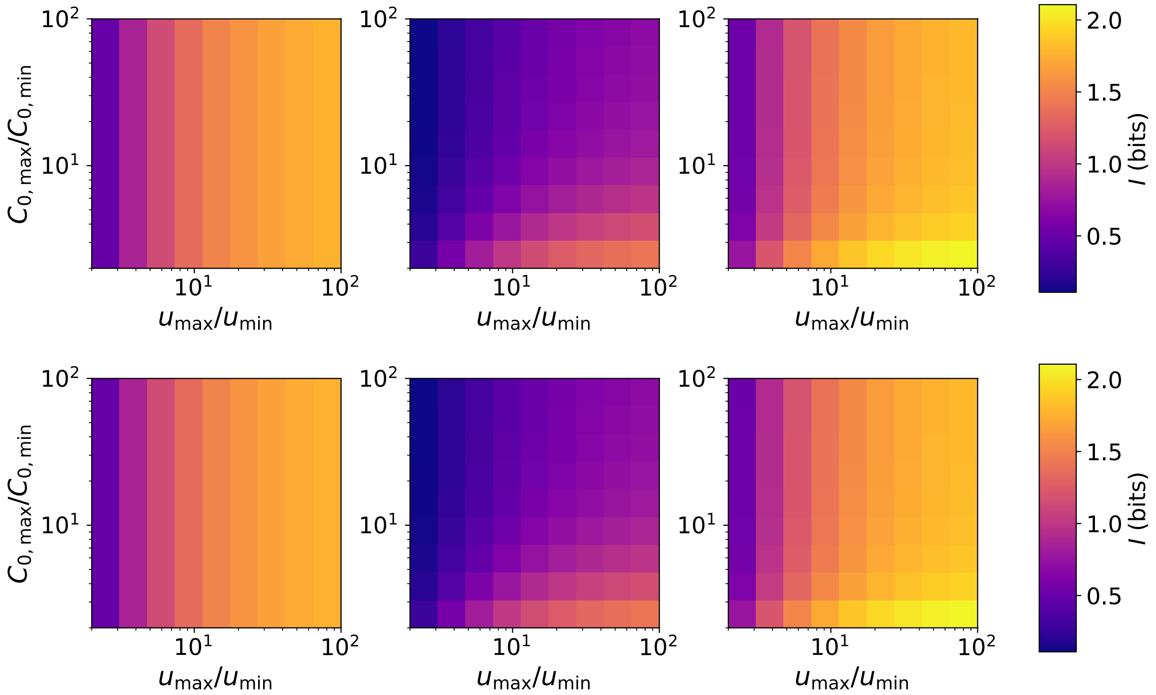


Supplementary Figure S5: Heat maps showing mutual information for different ranges of the uniform prior distribution when $u$ is the quantity of interest. The top row is for $u_{\rm min} = 10^2$ and bottom row is for $u_{\rm min} = 10^4$. The left, middle and right columns correspond to the unfolded protein-mediated, chaperon-mediated and hybrid mechanisms, respecively. The data points are calculated using $N = 100$.

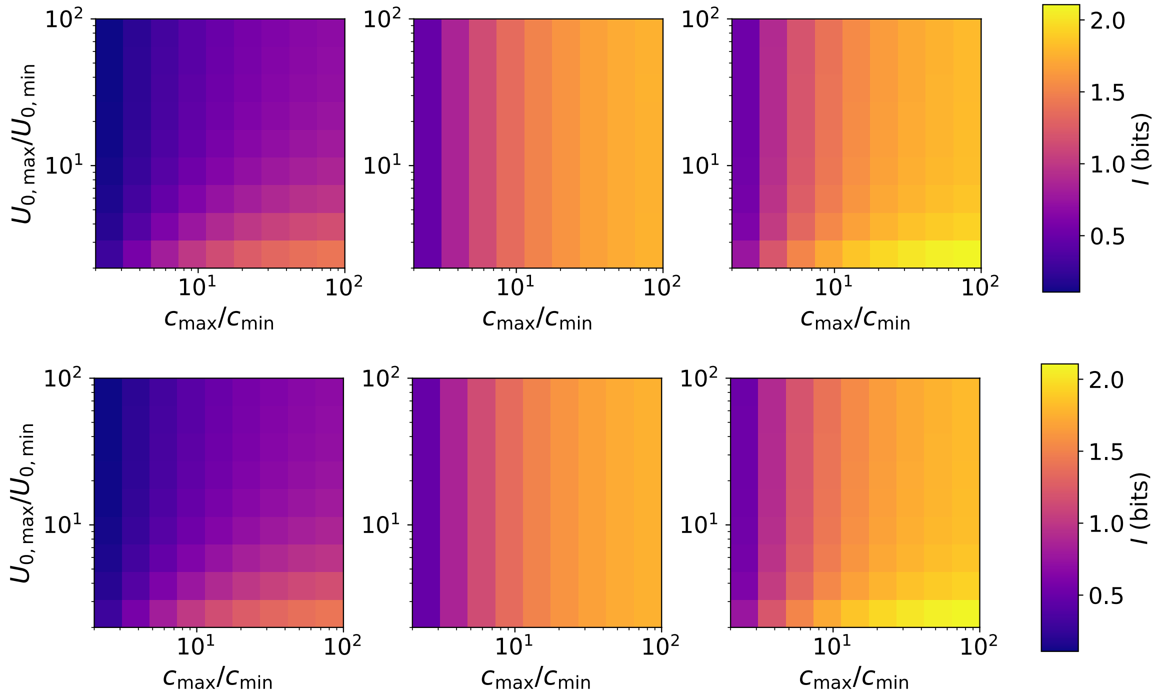


Supplementary Figure S6: Heat maps showing mutual information for different ranges of the uniform prior distribution when $c$ is the quantity of interest. The top row is for $c_{\mathrm{min}} = 10^2$ and bottom row is for $c_{\mathrm{min}} = 10^4$. The left, middle and right columns correspond to the unfolded protein-mediated, chaperon-mediated and hybrid mechanisms, respecively. The data points are calculated using $N = 100$.
